# Glucose-lactate metabolic cooperation in cancer: insights from a spatial mathematical model and implications for targeted therapy

**DOI:** 10.1101/008839

**Authors:** Jessica B. McGillen, Catherine J. Kelly, Alicia Martíez-González, Natasha K. Martin, Eamonn A. Gaffney, Philip K. Maini, Vıctor M. Pérez-García

## Abstract

A recent hypothesis has proposed a glucose-lactate metabolic symbiosis between adjacent hypoxic and oxygenated regions of a developing tumour, and proposed a treatment strategy to target this symbiosis. However, *in vivo* experimental support remains inconclusive. Here we develop a minimal spatial mathematical model of glucose-lactate metabolism to examine, in principle, whether metabolic symbiosis is plausible in human tumours, and to assess the potential impact of inhibiting it. We find that symbiosis is a robust feature of our model system—although on the length scale at which oxygen supply is diffusion-limited, its occurrence requires very high cellular metabolic activity—and that necrosis in the tumour core is reduced in the presence of symbiosis. Upon simulating therapeutic inhibition of lactate uptake, we predict that targeted treatment increases the extent of tissue oxygenation without increasing core necrosis. The oxygenation effect is correlated strongly with the extent of wildtype hypoxia and only weakly with wildtype symbiotic behaviour, and therefore may be promising for radiosensitisation of hypoxic, lactate-consuming tumours even if they do not exhibit a spatially well-defined symbiosis. Finally, we conduct a set of *in vitro* experiments on the U87 glioblastoma cell line to facilitate preliminary speculation as to where highly malignant tumours might fall in our parameter space, and find that these experiments suggest a weakly symbiotic regime for U87 cells, which raises the new question of what relationship exists between symbiosis—if indeed it occurs *in vivo*—and tumour malignancy.

## 1. Introduction

Recent years have seen a refinement of the historical view of tumours as greedy consumers of glucose. Experiments by Cheeti et al. (2006), Pavlides et al. (2009, 2010), and Sonveaux et al. (2008) indicate that tumour cells can consume lactate oxidatively as a metabolic substrate, either as a replacement fuel for glucose or concurrently with it. This lactate consumption has been shown to occur via oxidative phosphorylation (OXPHOS) and to depend upon uptake through the monocarboxylate transporter MCT1 (Sonveaux et al. 2008). On the basis of these discoveries, Sonveaux et al. (2008) put forward the hypothesis that “…lactate, the end-product of glycolysis, is the keystone of an exquisite symbiosis in which glycolytic and oxidative tumour cells mutually regulate their access to energy metabolites” in a spatially compartmentalised manner dependent upon oxygenation of the tumour tissue. Additionally, Sonveaux et al. (2008) proposed a novel treatment strategy, which centred on radio-sensitising a tumour by inhibiting MCT1 to force oxygenated cells to consume glucose and thereby starving nearby hypoxic, radiation-invulnerable, cells. Figure 1 provides a graphical interpretation of these ideas.

**Figure 1:**
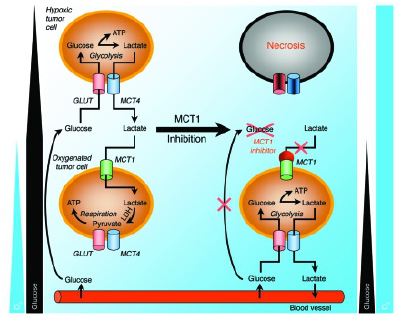
Metabolic symbiosis as hypothesised by Sonveaux et al. (2008). In a wildtype tumour (left panel), cells in the oxygenated environment near a capillary take up lactate for consumption by oxidative phosphorylation, gated by monocarboxylate transporter 1 (MCT1), while cells in the hypoxic environment of the tumour interior take up glucose for consumption by (anaerobic) glycolysis, gated by glucose transporters (GLUT) and producing lactate, which is then extruded by monocarboxylate transporter 4 (MCT4). This cooperation is thought to preserve glucose for the tumour interior. Sonveaux et al. postulated that inhibiting MCT1 (right panel) would force cells near the capillary to consume glucose instead of lactate, starving interior cells and increasing oxygenation into the tumour. Image reproduced with permission from the *Journal of Clinical Investigation*.

Sonveaux et al. (2008) presented elegant *in vitro* evidence for dual glucose-lactate metabolism gated by MCT1 in oxidative SiHa versus glycolytic WiDr tumour cells. Qualitative *in vivo* experiments on mouse LLc and human WiDr tumours further suggested that symbiosis was a robust tumour feature, and both the necrotic area and tissue oxygenation were shown to be extended by chronic MCT1 inhibition *in vivo*. However, a subsequent set of independent *in vivo* experiments carried out by Busk et al. (2011) on SiHa and FaDu_dd_ tumours saw no impact by MCT1 inhibition on either necrosis or the tissue glucose concentration, and only a transient decrease in the correlation between hypoxia and glucose uptake. Controversy has thus arisen as to whether the symbiosis hypothesis is an important feature of tumours and, especially, whether MCT1 inhibition of this symbiosis can cause the desired detrimental effects. he symbiosis hypothesis can be framed as a mathematical reaction-diffusion problem—how do the nutrients of interest flow in tumour tissue, and what spatial features develop as a consequence of consumption of these nutrients across the tumour domain?—and hence there is scope for mathematics to help resolve some of this controversy. By translating the dual glucose-lactate metabolic system into a mathematical model, with underlying assumptions made explicit, it may be possible to shed light on whether *in vivo* symbiosis as proposed by Sonveaux et al. (2008) is plausible in principle, and which of the two sets of experimental results is more likely to represent clinical reality.

Mendoza-Juez et al. (2012) developed a preliminary mathematical description of dual glucose-lactate tumour metabolism *in vitro*. This model was non-spatial and captured the metabolic behaviour of two fractional populations of tumour cells, one oxidative and the other glycolytic, in the presence of glucose and lactate. In brief, Mendoza-Juez et al. (2012) validated their model against five human cancer cell lines with varying characteristic metabolic rates and successfully replicated behaviours observed *in vitro* by Elstrom et al. (2004), Sonveaux et al. (2008), and Voisin et al. (2010). Their study thus supports the idea that tumour cells can consume lactate and that this capability, conferred by MCT1 expression, varies among cell lines in an *in vitro* environment. However, it leaves unanswered the question of whether, in principle, metabolic symbiosis is plausible *in vivo*.

The symbiosis hypothesis is intrinsically related to spatial tumour features, as the metabolism of each cell is responsive to hypoxia cues in the local microenvironment and hence based, at least in part, on relative distance from the capillary bed. Consequently, to fully capture the symbiosis hypothesis, a spatial model is needed. In this paper we develop a minimal spatial model of glucose-lactate metabolism and use it to interrogate both the plausibility of metabolic symbiosis *in vivo* and the potential effectiveness of blocking this symbiosis as a radiosensitisation strategy. We find that symbiosis is a robust feature of our model system, and, although it does not directly rescue tumours from necrosis, on average its effects on the tumour dynamics are in line with expectations. However, symbiosis does require the cells’ metabolic activity to be several orders of magnitude higher than what is commonly observed *in vitro*. Whether such levels are plausible remains an open question. Furthermore, we simulate MCT1 inhibition and discover that, in agreement with Busk et al. (2011), this blocking of symbiosis does not cause the expected increase in necrosis; but, in agreement with Sonveaux et al. (2008), it does cause an increase in tissue oxygenation. This oxygenation is highly correlated in our model with the extent of hypoxia exhibited by the wildtype system, suggesting that MCT1 inhibition may be a promising strategy for radiosensitising lactate-consuming tumours with or without symbiosis. Finally, we demonstrate experimentally that U87 glioblastoma tumours are likely to fall in a parameter regime that produces only weakly symbiotic behaviour, raising an open question of how symbiosis relates to malignancy in general.

## 2. Our model

Throughout this study, we will aim for a model of minimal complexity, conjecturing that if metabolic symbiosis is not possible in a simplified scenario where glucose and lactate dominate the metabolic landscape, then it is unlikely to be a significant feature in more complicated scenarios. We do consider it vital for our model to be spatial, however, as the metabolic symbiosis hypothesis *in vivo* is a spatial feature; hence, a minimal system of partial differential equations provides the requisite level of complexity.

### 2.1 Motivation and framework

The model we develop here is a direct extension, with simplifications, of the model proposed by Mendoza-Juez et al. (2012) to a spatial tumour scenario. As such, it captures behaviours that are qualitatively similar to those with which the non-spatial model was verified, namely patterns of dual consumption of glucose and lactate that vary across tumour cell lines depending on the relative expression of membrane transporters. However, we simplify the modelling framework considerably by avoiding the assumption of metabolic switching, and instead suppose that cells can use both glycolysis and oxidative phosphorylation given their relative expression of the requisite membrane transporters and the appropriate environment conditions (e.g. oxidative phosphorylation (OXPHOS) requires oxygen). This change allows us to avoid imposing an artificial dichotomy between the metabolic pathways. We depart further from the Mendoza-Juez et al. (2012) model by neglecting consumption of glucose by oxidative phosphorylation, which experiments have shown to be a minor factor relative to glycolysis in proliferative cells (Pfeiffer et al. 2001, Vander Heiden et al. 2009, Koppenol et al. 2011). An alternative model that does consider OXPHOS of glucose can be found in the Supplemental Information and exhibits dynamics which are qualitatively similar to those presented here.

We model a one-dimensional ray in polar coordinates extending out from the core of a tumour, under the assumption of spherical symmetry, to a spherical shell of capillaries 0.02 cm from the core. This modelling scenario corresponds either to a small avascular tumour or, more interestingly (and more realistically biologically), to pockets of tissue inside a larger vascularised tumour. Large tumours exhibit heterogeneities in oxygen supply due to capillary crushing and leakiness of tumour-induced vascular networks (Gatenby et al. 2007, Gillies & Gatenby 2007, Basanta et al. 2011), and hence our model can be viewed as a simplified representation of localised metabolic features in a heterogeneous tumour.

### 2.2 Equations and terms

The species in our model are dependent upon time, *t*, in days, and one-dimensional spatial coordinate, *r*, in centimetres from the tumour core, or interior of the local tumour pocket. These species are cell density, *C*(*r, t*), normalised by the tissue carrying capacity; the tissue concentration of glucose, *G*(*r, t*), in mM; the tissue concentration of lactate, *L*(*r, t*), in mM; and the partial pressure of oxygen, *O*(*r, t*), in mM. We model only tumour tissue as inclusion of the underlying stroma could introduce tumour-healthy metabolic interactions (Pavlides et al. 2009, 2010), and we wish to keep the modelling scenario restricted to a simple in-principle test of the symbiosis hypothesis as it was stated by Sonveaux et al. (2008).

We let *ρ* represent the rate of tumour cell growth; *δ* the rate of tumour cell necrosis; and *κ_G_* and *κ_L_* the maximal rates of consumption of glucose and lactate, respectively, by the tumour population. Additionally defining switch functions capturing tumour sensitivity to hypoxia, glucose deprivation, severe acidosis, and severe anoxia as *ψ_H_*, *ψ_G_*, *ψ_A_*, and *ψ_N_*, respectively, our model equations are

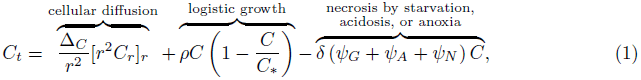

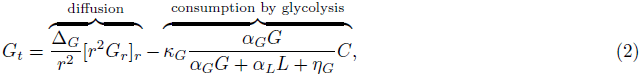

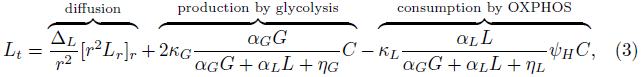

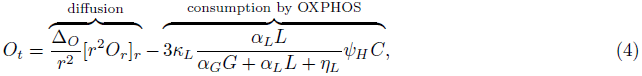

where Δ*_C_*, Δ*_G_*, Δ*_L_*, and Δ*_O_* denote the diffusion coefficients of tumour cells, glucose, lactate, and oxygen, respectively. We assume a Michaelis-Menten form for the metabolite uptake functions such that *η_G_* and *η_L_* represent the half-saturation points for uptake of glucose and lactate, respectively.

Although expression of metabolite transporters is tied to the microenvironment *in vivo*, with GLUT expression being hypoxia-induced (Burgman et al. 2001, Airley et al. 2001, Williams et al. 2002, Wincewicz et al. 2007, Liu et al. 2009) and MCT1 expression being hypoxia-repressed (Berra et al. 2003, Marxsen et al. 2004, Sonveaux et al. 2008, Feron 2009, Semenza 2010), different cell lines exhibit different characteristic levels of expression *in vitro* (Elstrom et al. 2004, Sonveaux et al. 2008, Voisin et al. 2010) and it is possible that cells retain these features *in vivo* in the form of characteristic maximal levels of expression. The parameters *α_G_* and *α_L_* in Equations (2)–(4) allow for this possibility: if the cells’ characteristic capacity for MCT1 expression is high relative to that for GLUT, then we will have *α_L_ > α_G_*; conversely, if the characteristic capacity for MCT1 expression is low relative to that for GLUT, then we will have *α_L_ < α_G_*. If *α_G_* = *α_L_* then the cells have no characteristic capacities *in vivo* and instead transporter expression is determined entirely by the microenvironment. We note that Equations (2)–(4) preserve the appropriate stoichiometry of the metabolic pathways; that is, two molecules of lactate are produced per molecule of glucose consumed by the glycolytic pathway and three molecules of oxygen are consumed per molecule of lactate processed by OXPHOS.

Tumour cells may become necrotic under severely glucose-starved, acidic, or anoxic conditions. We model this in Equation (1) as a sum of simple sensitivity switches—*ψ_G_* for glucose starvation, *ψ_A_* for acidosis, and *ψ_N_* for anoxia—such that the speed of cell death increases as conditions become harsher. Additionally, OXPHOS is sensitive to hypoxia, which we represent as an oxygen-dependent switch, *ψ_H_*, multiplying the uptake of lactate in Equations (3) and (4). These four sensitivity switches are

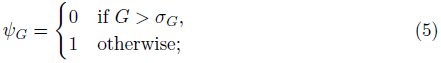

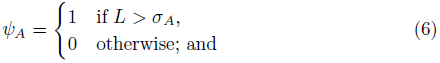

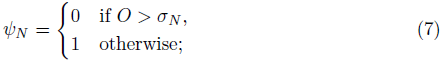

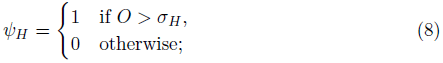

where *σ_G_* denotes the minimum glucose concentration that cells require for survival, *σ_A_* the lactate concentration corresponding to the maximum tolerable level of tissue acidity, *σ_N_* the oxygen threshold for severe anoxia, and *σ_H_* the oxygen threshold for hypoxia. We note that *σ_G_* and *σ_A_* are difficult to measure experimentally, and hence these constants are somewhat abstract. Nevertheless, we will make the reasonable choices of setting *σ_G_* to be small and *σ_A_* to be above the lactate level typically observed in tumour tissue (Herholz et al. 1992).

In keeping with Mendoza-Juez et al. (2012), we make the slight simplification of scaling the tumour cell density by the tissue carrying capacity, such that C̃ = *C/C*_∗_. Further letting *a* = *α_L_/α_G_*, *n_G_* = *η_G_/α_G_*, *n_L_* = *η_L_/α_L_*, *k_G_* = *κ_G_C_∗_*, and *k_L_* = *κ_L_C_∗_*, and dropping the tilde for convenience, we have

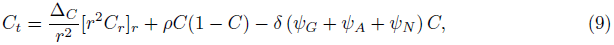

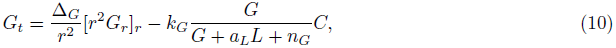

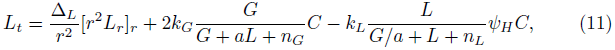

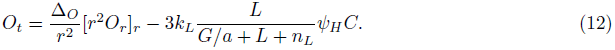

Lastly, for later notational convenience we define the uptake functions

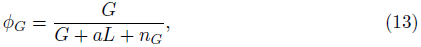

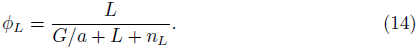

Boundary conditions are zero-flux for all species in the tumour core, under our assumption of spherical symmetry, and we retain this zero-flux condition for the tumour population at the edge of the tumour (at *r* = *R* where *R* is the radius of a typical avascular tumour, taken here to be 0.02 cm), such that the cells cannot migrate beyond the capillary shell. Metabolites are permitted to exchange with the capillary shell as follows (for a generic metabolite, *M)*:

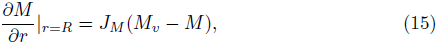

where *J_M_* is the coefficient of exchange and *M_v_* the concentration of the metabolite in the vessels. The vessels act as a source for glucose and oxygen and, commonly, as a sink for lactate. In exceptional cases of chronic hyperlactatemia (Huckabee 1961, Levraut et al. 1998, Boubaker et al. 2001) or insulin-induced hypoglycemia (Boumezbeur et al. 2010), the capillary shell may act instead as a source for lactate, but we do not consider this relatively rare phenomenon here. The vessel concentrations of glucose, lactate, and oxygen (*G_v_*, *L_v_*, and *O_v_*, respectively) and corresponding exchange coefficients (*J_G_*, *J_L_*, and *J_O_*) can be found in Table 1.

**Table 1:** Parameters in Equations (9)–(12). Model parameters (with dimensions, as we have only partially scaled the system): listed are value estimates for parameters that we consider fixed in order to set up a consistent background against which to view the dynamics of tumour glucose-lactate consumption (top portion), and ranges for our metabolic parameters that are of interest as varying in a meaningful way across tumours (bottom portion)

|  | Parameter | Value | Units | Source |
| --- | --- | --- | --- | --- |
| Fixed | $\Delta_C$ | $5.7 \cdot 10^{-6}$ | cm<sup>2</sup>/day | Rockne et al. (2010) |
| | $\Delta_G$ | 0.0346 | cm<sup>2</sup>/day | Jiang et al. (2005) |
| | $\Delta_L$ | 0.0518 | cm<sup>2</sup>/day | $1.5\Delta_G$, due to size |
| | $\Delta_O$ | 0.864 | cm<sup>2</sup>/day | Rockne et al. (2010) |
| | $\rho$ | 1/17 | 1/day | Rockne et al. (2010) |
| | $\delta$ | 1/10 | 1/day | Estimated |
| | $\sigma_H$ | 0.0278 | mM | Daşu et al. (2003), converted from mmHg using Henry's Law |
| | $\sigma_G$ | 0.1 | mM | Chosen to be small |
| | $\sigma_A$ | 20 | mM | Chosen to be large |
| | $\sigma_N$ | 0.001 | mM | Chosen to be small |
| | $J_G$ | $10^3$ | 1/cm | High as diffusion across vessel wall is facilitated (McEwen & Reagan 2004) |
| | $J_O$ | $10^3$ | 1/cm | High as the capillary wall offers negligible resistance to oxygen diffusion (Blum 1960) |
| | $J_L$ | 10 | 1/cm | Assumed moderate for functioning vessels |
| | $G_v$ | 5 | mM | Physiological (Sherwood 2007) |
| | $L_v$ | 0 | mM | Physiological (Figley 2011) |
| | $O_v$ | 0.0834 | mM | Physiological (Sherwood 2007), converted from mmHg using Henry's Law |
| Varied | $k_G$ | $[1, 10^5]$ | mM/day | Chosen to cover a large range |
| | $k_L$ | $[1, 10^5]$ | mM/day | Chosen to cover a large range |
| | $n_L$ | $[0.01, 10]$ | mM | Mendoza-Juez et al. (2012) |
| | $n_G$ | $[0.01, 10]$ | mM | Mendoza-Juez et al. (2012) |
| | $a$ | $[10^{-3}, 10^3]$ | none | Chosen to cover a large range |

We impose initial conditions such that the tumour cells have a uniform density of half the carrying capacity across the domain, and metabolite concentrations take a uniform value equal to their vessel concentrations. The steady-state results of our system are qualitatively insensitive to our choice of initial conditions, and we have chosen this particular configuration to facilitate a clear interpretation of any necrosis in the tumour core that we may see relative to the edge.

### 2.3 Metrics for model exploration

‘Symbiosis’ as proposed by Sonveaux et al. (2008) is a qualitative and imprecisely defined feature of the tumour system. Nevertheless, we can consider it by determining the fraction of total metabolic consumption occupied by glycolysis, denoted Φ*_g_*, and the fraction occupied by oxidative phosphorylation of lactate, denoted Φ*_l_*; these are given by:

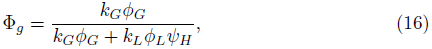

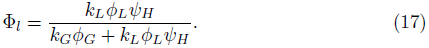

We consider symbiosis to occur when glycolysis dominates over oxidative phosphorylation of lactate (OXPHOS) in the hypoxic core, and OXPHOS dominates over glycolysis in the oxygenated region near the tumour edge. Further, we can consider the ‘strength’ of symbiosis to be given by the degree of dominance OXPHOS takes over glycolysis in the oxidative region. We are also interested in the occurrence of necrosis, or a reduction in tumour density in the core relative to the edge, and the extent of hypoxia, or the location along the tumour domain where oxygen crosses the hypoxia threshold, *σ_H_*. Accordingly, letting *θ* denote symbiosis, *γ* denote necrosis, and *λ* denote the extent of hypoxia, we define the metrics

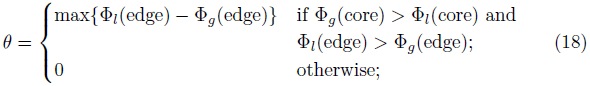

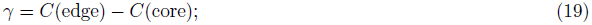

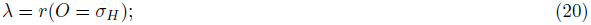

with *λ* subject to the limiting cases that *λ* = 0 if the whole tissue is normoxic and *λ* = *R* if the whole tissue is hypoxic.

## 3. Computational and experimental methods

### 3.1 Numerical solutions

Throughout this chapter, Equations (9)–(12) are solved numerically using the Method of Lines (Schiesser & Griffiths 2009). By this method, we discretise space using a very fine step (*dr* = 10*^−^*^4^ cm) and solve the resulting system of coupled ordinary differential equations through time using Matlab’s inbuilt ODE15s solver to accommodate stiffness.

Numerically simulating Equations (9)–(12) with a representative choice of parameters reveals that an end-time of 200 days is sufficient to achieve a steady state in which all species have reached a plateau (Figure 2).

**Figure 2:**
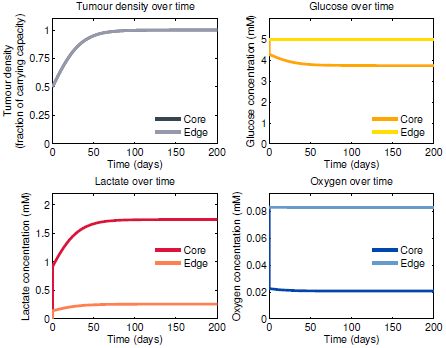
Temporal behaviour of Equations (9)–(12). Over-time profiles in the tumour core (darker colours) and at the tumour edge (lighter colours) for the tumour cell density (greys) and concentrations of glucose (oranges), lactate (reds), and oxygen (blues), for a representative choice of parameters: *a* = 1, *n_L_* = 1, *n_G_* = 1, *k_L_* = 10^4^, and *k_G_* = 10^3^, with the remaining parameters as listed in the top portion of Table 1. All profiles reach steady state well before the simulation end-time of 200 days.

For the remainder of this work, we will be concerned with spatial behaviours established at steady state. Hence, all simulations are run to an end-time of 200 days; parameters are listed in Table 1 and discussed further in Section 3.2.

In Figure 2 it appears that we have fast early-time dynamics, possibly due to the magnitude of the uptake rate parameters *k_G_* and *k_L_*. A full nondimension-alisation could help to elucidate the relative timescales driving the system, and an asymptotic analysis could potentially generate insight. However, we wish to explore the full range of possible parameter values for our model—including smaller choices for *k_L_* and *k_G_*—in order to allow scope for metabolic aberrations, such as mitochondrial dysregulation, which are observed in many tumours and which perturb the normal stoichiometry of ATP production (Seyfried & Shelton 2010). We therefore turn our attention to an exploration of some effective numerical methods by which to obtain a comprehensive picture of the fundamental parameter space of the model. Such methods can serve as a kind of proxy for analytical insight in differential equation settings and have an elegance and power of their own.

### 3.2 Multivariate sensitivity analysis

In any given model of a biological system, some of the fundamental model parameters may be well-determined, but others either cannot be measured accurately due to experimental limitations, or are of interest where variation is an important biological feature—for example, across different tumours. To explore the sensitivity of a system to its parameters that fall into the latter category, i.e. are of interest as having the potential to vary, it is possible to carry out a standard single-parameter or local sensitivity analysis in which one parameter is varied at a time while the others are held constant at ‘baseline’ or reference values (Marino et al. 2008). However, such an approach, while common, loses information about underlying uncertainties—for example in estimation of the ‘baseline’ values—and also about parameter interaction effects, and so would provide incomplete insight into the system (James & McCulloch 1990, Marino et al. 2008, Saltelli & Annoni 2010).

To escape the limitations of one-at-a-time sensitivity, we can simply allow the parameters of interest to vary simultaneously, and take random samples from the resulting multi-dimensional parameter space (Saltelli et al. 2008) for use in repeated numerical solutions of the model system. We can then examine the resulting distribution of model behaviours using histograms, scatterplots, and further statistical methods as required. While more sophisticated approaches to global sensitivity analysis exist (Wu et al. 2013), for our system a sampling-based approach is sufficient to avoid the pitfalls of single-parameter sensitivity analyses while remaining computationally tractable.

The parameters in Equations (9)–(12) which we fix and consider ‘background’ parameters in order to set up a consistent “background” system against which to view the tumour metabolic dynamics, are listed with their sources in the top portion of Table 1. The remaining five parameters, which are the metabolic parameters and therefore of interest as potentially varying across tumours, are sampled from the ranges listed in the lower portion of Table 1, for repeated numerical solutions of Equations (9)–(12) to steady state as described in Section 3.1.

Global, multidimensional sensitivity analyses have been gaining traction in systems biology (Marino et al. 2008) and nonlinear modelling (Homma & Saltelli 1996), particularly of ecology (De’ath & Fabricius 2000, Fieberg & Jenkins 2005, Cutler et al. 2007, Cariboni et al. 2007) and infectious diseases (Wu et al. 2013). Yet, to our knowledge, we are the first to apply such approaches to tumour metabolism modelling. In the following we briefly discuss two complementary algorithms—one linear and one nonlinear—that exploit this multidimensional, sampling-based method for sensitivity analysis.

#### 3.2.1 Principal component analysis

Principal component analysis (PCA) is a simple, non-parametric approach for extracting relevant information from large, complex datasets by reducing their dimensionality (Hastie et al. 2009). The PCA transformation is defined such that the first principal component captures as much variability in the data as possible, and each subsequent component captures the largest possible amount of the remaining variability, subject to the constraint that the new component is uncorrelated with (i.e. orthogonal to) the preceding components.

If a dataset **X** is arranged as an *m*-by-*n* matrix such that each of *n* observations comprises a column vector of *m* variables (here, parameters), then each observation (here, a sampled point from our model parameter space) is a vector that lies in an *m*-dimensional vector space spanned by a basis. The goal of PCA is to find an alternative basis which is a linear combination of the original basis vectors and expresses the dataset more meaningfully. Note that PCA is a linear method; as such, it is stringent, because most complex systems are nonlinear, but also powerful, because it renders the problem amenable to linear algebra.

Letting **X** and **Y** denote *m*-by-*n* matrices related by a linear transformation **P**, such that **X** is the original dataset and **Y** is a re-representation of it, and further defining 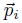 as the rows of **P**, 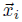 as the columns of **X**, and 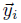 as the columns of **Y**, a change of basis is represented by the equation **PX** = **Y**. The rows of **P**, 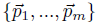 are a set of new basis vectors for expressing the columns of **X**: by construction, the principal components.

The covariance matrix for the dataset **X** is then given by 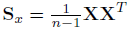, a square symmetric *m*-by-*m* matrix. As our goal is to capture meaningful features of the data, we want to remove the redundancy (i.e. covariance) from the variables, which, in turn, means diagonalising **XX***^T^*. This operation ensures that all basis vectors 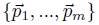 orthonormal, and carries the assumption that the directions with the largest variances are the most important, i.e. principal.

#### 3.2.2 Analysis of parameter importance using classification trees

We can explore the sensitivity of a model output of interest to the fundamental model parameters—or, equivalently, the relative importance of the parameters for classifying that output—by applying a tree-based statistical classifier (Pappenberger et al. 2006). A system of partial differential equations can be translated into a classification problem by first generating a large random sample of the multidimensional parameter space, then numerically simulating the system of partial differential equations to steady state and applying the binary classification

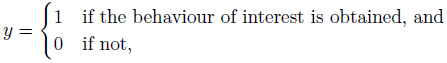

for each of the sampled parameter points. Our model parameters subsequently comprise the set of variables for a tree analysis and our vector *y* the classified responses. Tree-based methods are nonlinear and iteratively determine which variable best classifies the training data, progressively segmenting the data by further classifications as the tree grows deeper.

Trees can capture complex interaction structures in data, but also are inherently noisy (Hastie et al. 2009), as a small error near the top may propagate into a large effect at the bottom. A remedy for this is the popular machine-learning algorithm RandomForest, developed by Breiman (2001), which grows a forest of trees, each on an independent bootstrapped sample (random sample with replacement) from the training data, and averages over the predictions from all trees to reduce the variance in the final result. RandomForest is fast, accurate, and capable of handling a high-dimensional variable space.

Each tree in a given forest has a built-in test set—the out-of-bag data, or data not included in the bootstrapped sample for that tree—which can be exploited to determine the relative importance of the variables for accurate classification. After growing a forest, the class of each point in the training data is predicted using each tree for which that point is out-of-bag. The values of a variable *x* are then randomly permuted while holding the others fixed, and the permuted data are passed down the tree. The relative importance of variable *x* for the behaviour of interest is the difference in out-of-bag error before and after the permutation, averaged over all trees (Breiman 2001).

### 3.3. In vitro experiments

Sonveaux et al. (2008) and Busk et al. (2011) considered the highly oxidative cervical cell line SiHa, the glycolytic colon line WiDr, and the glycolytic squamous carcinoma line FaDu_dd_ *in vitro* and *in vivo*. Mendoza-Juez et al. (2012) additionally considered results from *in vitro* experiments on two glycolytic glioma lines (LN18 and LN229) and a more oxidative glioma line (C6). We supplement these data further with a set of *in vitro* experiments carried out on the highly malignant glioblastoma cell line U87. This extends our modelling scenario to glioblastoma tumours, which are notoriously challenging clinically and hence of special interest. Moreover, the blood-brain barrier renders glioma tumours slightly metabolically simplified—at least in terms of the variety of metabolic substrates present in the local tissue—in comparison to body tumours (Terpstra et al. 1998, Moreno-Sánchez et al. 2007), and thus U87 may be a good target for future model validation in vivo. Finally, considering this cell line opens the possibility for an eventual connection of tumour metabolic dynamics with healthy metabolic cooperation that is known to occur between glutamatergic neurons and glia (Kirchhoff et al. 2001, Pellerin & Magistretti 2004, Gladden 2004, Magistretti 2006, Pellerin et al. 2007, Bélanger et al. 2011).

We carried out a set of experiments to explore the metabolic characteristics of U87 glioblastoma cultures *in vitro* using three metrics: glucose uptake, extracellular acidification, and oxygen consumption. Glucose was measured using a radiolabelled analogue, [18F] fluorodeoxyglucose (FDG), which has similar uptake kinetics to glucose. Unlike glucose, FDG is trapped in cells, and is frequently used as an estimator of glycolytic flux. The rate of media acidification (i.e. the change in pH over time) provides an estimate for the rate of lactate production and additional marker of glycolysis. The rate of oxygen consumption reflects the rate of oxidative phosphorylation.

#### 3.3.1. Acidification and oxygen consumption

In a pair of experiments, U87 cells were seeded in a Seahorse XF Analyser 96-well cellplate (Seahorse Biosciences, USA) and grown in standard tissue culture under high-glucose conditions (25 mM) for 48 hours to allow full adhesion. For the assay, cells were incubated with 180 µl Seahorse medium (5 mM glucose, 4 mM glutamine, and 0.1 mM sodium pyruvate) to replicate physiological conditions. Basal rates of oxygen consumption and extracellular acidification were measured using the Seahorse XF Analyser. Sodium lactate was then injected at concentrations of 0, 3, 6, 9, 12, and 15 mM and further rate data were collected. Rate data were normalised to cell density as determined using Hoechst fluorescence staining.

#### 3.3.2. Uptake of glucose

U87 cells were trypsinised to a single cell suspension and transferred to counting tubes. Cells were washed twice with PBS and the growth medium was replaced with a medium containing 100 kBq [18F]FDG, 4 mM glutamine, and sodium lactate at concentrations of 0, 3, 6, 9, 12, and 15 mM. Cells were incubated for one hour at 37C; thereafter, cells were washed in PBS and activity in the cells was counted using a Beckman Coulter counter.

We note that these experiments consider only low to moderate lactate concentrations. Although lactate has been estimated to potentially reach as high as 35 mM in human brain tumours, maximum accumulation tends to occur in necrotic and cystic regions, with concentrations remaining below 15 mM in the more peripheral regions (Herholz et al. 1992) which are of interest for the symbiosis hypothesis.

## 4. Results

Application of the complementary statistical techniques for multivariate sensitivity analysis that we discussed in 3.2—importance analysis of symbiosis (*θ*), necrosis (*γ*), and hypoxia (*λ*) using classification trees, and principal component analysis of symbiosis (*θ*)—enables us to elucidate features of our multidimensional parameter space, as follows.

### 4.1. Consumption rates and substrate preference govern symbiosis

Our principal component analysis, a linear method, shows that the first two principal components for symbiosis (*θ*) comprise mainly the rates of glucose and lactate consumption by the tumour—*k_G_* and *k_L_*, respectively—and the characteristic preference for lactate uptake, *a* (Figure 3a). Our complementary, nonlinear, importance analysis is in agreement with this result (Figure 3b), suggesting that we are capturing robust dependencies.

**Figure 3:**
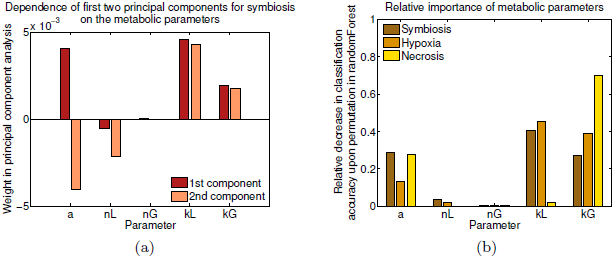
Principal component weighting and importance of parameters. (a) Weights assigned to each parameter from the original parameter space of Equations (9)–(12) in linear combinations comprising the first two principal components for symbiosis, described in Section 3.2.1; and (b) relative importance, as defined in Section 3.2.2, of the parameters for classifying the presence of symbiosis, *θ* (brown), tissue hypoxia, *λ* (tan), and necrosis, *γ* (yellow). The parameters are: expression of MCT1 relative to GLUT (*a*), the half-saturation point for uptake of lactate (*n_L_*) and glucose (*n_G_*), and the rates of consumption of lactate (*k_L_*), and glucose (*k_G_*).

Figure 3b also reveals that the importance weighting for the extent of hypoxia is similar (though not identical) to that for symbiosis, with lactate consumption (*k_L_*) the most important parameter for both, followed by glucose consumption (*k_G_*) and then preference for lactate (*a*). However, the importance weighting for necrosis is markedly different from these (Figure 3b), with necrosis depending primarily on consumption of glucose (*k_G_*), secondarily on preference for lactate (*a*), and much less on consumption of lactate (*k_L_*).

This difference in parameter dependence is further illustrated by projecting each of the three characteristics—symbiosis, necrosis, and hypoxia—onto the principal component space for symbiosis (Figure 4). Symbiosis and hypoxia project similarly (though not identically) to one another, while necrosis projects differently from both.

**Figure 4:**
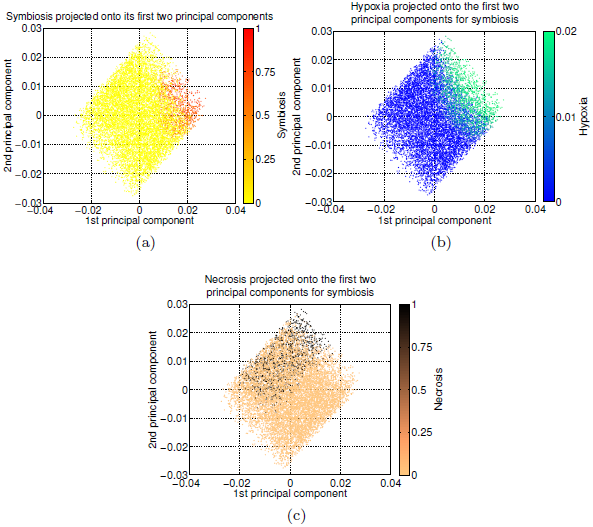
Projection onto the symbiosis principal component space. Projection onto the first two principal components for symbiosis, as defined in Section 3.2.1, of (a) symbiosis, *θ*, (b) hypoxia, *λ*, and (c) necrosis, *γ*. The projection of symbiosis onto the space of its first two principal components is similar to the projection of hypoxia onto the same space, while the projection of necrosis onto the same space is different from these.

The Michaelis-Menten constants for glucose and lactate uptake (*n_G_* and *n_L_*, respectively) have relatively small principal component weightings for symbiosis and are relatively unimportant for all three characteristics. This is perhaps surprising in view of the variability in *n_L_* across tumour cell lines *in vitro* (Mendoza-Juez et al. 2012).

### 4.2. Symbiosis exhibits the expected effects

Plotting symbiosis, necrosis, and hypoxia each against the three most important parameters pairwise (Figure 5) indicates that symbiosis can occur when consumption of glucose (*k_G_*) and of lactate (*k_L_*) are both high and the preference for lactate (*a*) is not small (top row of Figure 5). Additionally, broadly speaking it appears that symbiosis is more pronounced when *k_L_ > k_G_*. We note that symbiosis requires the tumour to be extremely active metabolically, as *k_G_* must be above 10^2^ mM/day and *k_L_* above 10^3^ mM/day for it to occur. Quantitative measurements of *in vivo* metabolic consumption are, to our knowledge, difficult to come by, but future experiments could potentially be focussed in this direction to help determine whether these are plausible parameter ranges for tumours *in vivo*.

**Figure 5:**
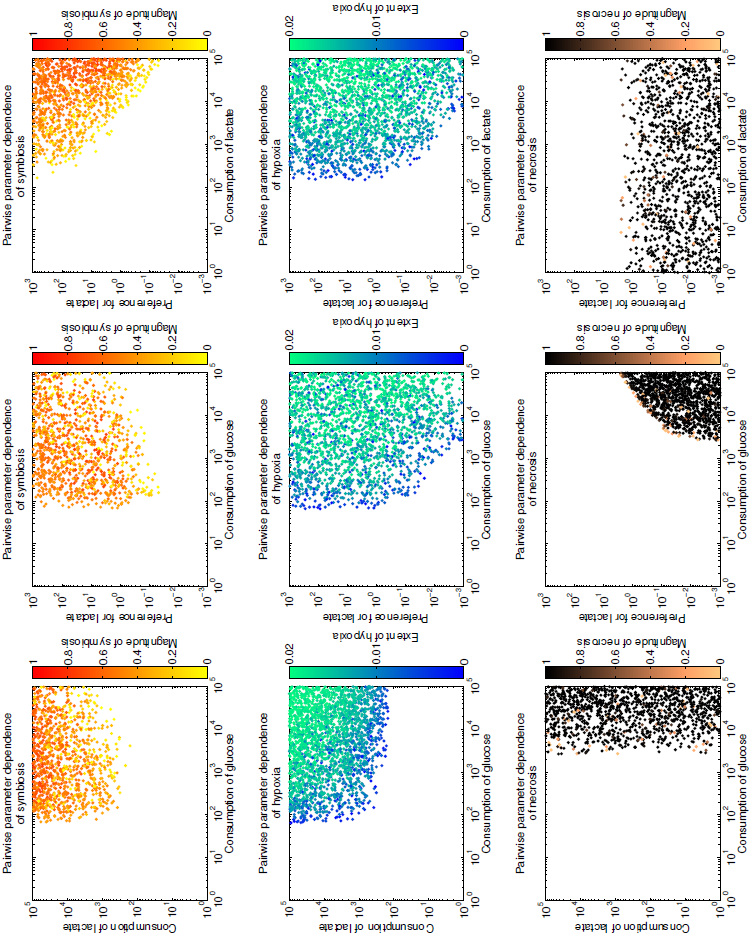
Symbiosis, hypoxia, and necrosis in the space of important parameters. Symbiosis (top row), hypoxia (middle row), and necrosis (bottom row), for numerical simulations of Equations (9)–(12) with 10^4^ random samples of the parameter space, plotted pairwise against the three most important parameters: rates of consumption of glucose (*k_G_*) and lactate (*k_L_*), and characteristic preference for lactate (*a*). (c)

The extent of hypoxia is highest when consumption of glucose and lactate (*k_G_* and *k_L_*, respectively) are both very high, though it is less dependent than symbiosis on the preference for lactate (*a*) (middle row of Figure 5). The fact that hypoxia does not occur in the regime of lower metabolic consumption rates suggests that metabolically less-active tumours of the size considered here do not exhibit the nutrient gradients that are needed to drive the development of a spatial symbiosis.

Necrosis is mostly—though not entirely—mutually exclusive to symbiosis, reaching high values when glucose consumption (*k_G_*) is very high and the preference for lactate (*a*) is small (bottom row of Figure 5). Plotting a representative sample from this region of parameter space over time reveals that the cause of this necrosis is glucose starvation, rather than acidosis or severe anoxia (Figure 6). There is, however, a small region of parameter space in which symbiosis and necrosis are not mutually exclusive: when the preference for lactate (*a*) is on the order of one, and consumption of glucose (*k_G_*) is on the order of 10^4^, both symbiosis and necrosis can occur. It would be informative to determine whether or not this small parameter region is narrow enough to be considered a biological fine-tuning; but this would require a mapping between this model parameter region and the space of clinically plausible tumour characteristics, which again points to the need for quantitative *in vivo* measurements of the metabolic parameters.

**Figure 6:**
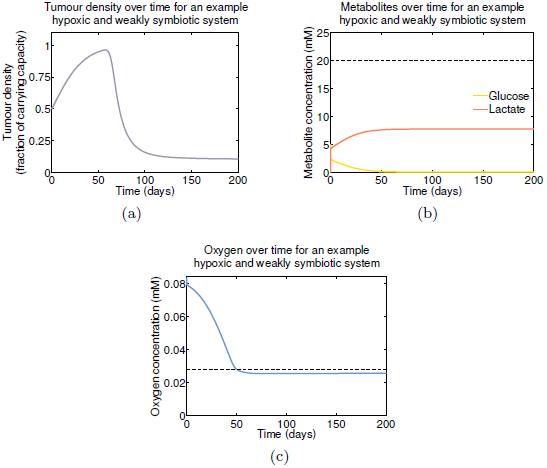
Cause of necrosis under weak/no symbiosis. Species profiles in the tumour core over time for a representative sample from the weakly-/non-symbiotic parameter regime governing Equations (9)–(12). Shown are the (a) tumour density (grey), (b) glucose (yellow) and lactate (red) concentrations, and (c) oxygen concentration (blue). Dotted lines in (b) and (c) indicate the acidosis threshold *σ_A_* and the hypoxia threshold *σ_H_,* respectively.

Figures 7 and 8 illustrate the averaged steady-state system behaviours over the tumour domain for the parameter regimes which produce either strongly symbiotic behaviour, with lactate consumption dominating over glucose consumption near the oxygenated tumour edge (Figure 7), or weakly-or non-symbiotic behaviour, with lactate consumption at the tumour edge which does not strikingly dominate over glucose consumption at the tumour edge (Figure 8). On average, the strongly-symbiotic regime exhibits lower lactate, higher glucose, and a greater extent of hypoxia over the tumour domain than does the less-symbiotic regime. The latter additionally exhibits considerable necrosis in the tumour core.

**Figure 7:**
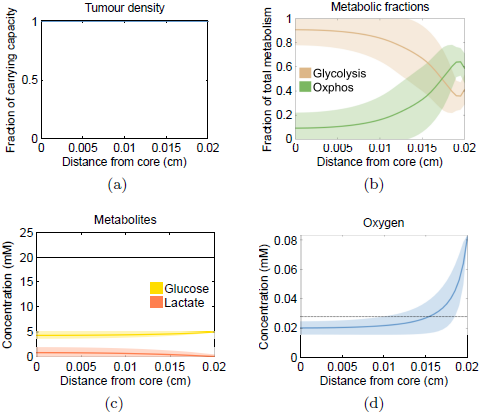
Symbiotic behaviour of Equations (9)–(12). Averaged spatial behaviours of the (a) tumour density, (b) metabolic fractions, (c) metabolite concentrations, and (d) oxygen concentration, at steady state. Shown are the means (curves) and standard deviations (shaded areas) from 10^4^ uniformly distributed samples from the parameter region giving rise to symbiotic behaviour; that is, from the region in which consumption of lactate (*k_L_*) and glucose (*k_G_*) are high with *k_L_ > k_G_*, and the characteristic preference for lactate (*a*) is high. Fixed parameters are listed in the top portion of Table 1.

**Figure 8:**
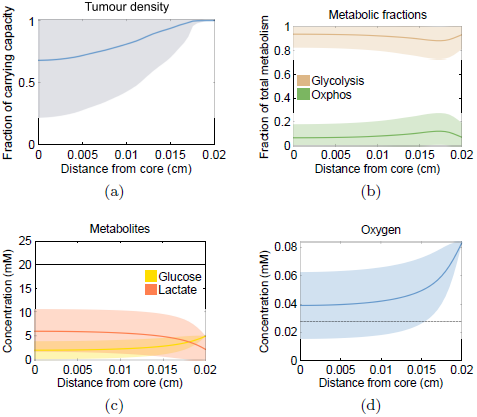
Weakly-or non-symbiotic behaviour of Equations (9)–(12). Averaged spatial behaviours of the (a) tumour density, (b) metabolic fractions, (c) metabolite concentrations, and (d) oxygen concentration, at steady state. Shown are the means (curves) and standard deviations (shaded areas) from 10^4^ uniformly distributed samples from the parameter region giving rise to weakly-or non-symbiotic behaviour; that is, from the region in which consumption of lactate (*k_L_*) and glucose (*k_G_*) are high with *k_G_ > k_L_*, and the characteristic preference for lactate (*a*) is low. Fixed parameters are listed in the top portion of Table 1.

### 4.3. MCT1 inhibition increases oxygenation but not necrosis

The first part of the symbiosis hypothesis concerns the establishment and beneficial effects of a well-defined spatial symbiosis, which we have considered thus far. The second part proposes that inhibiting symbiosis by blocking the monocarboxylate transporter 1 (MCT1) forces cells near the tumour edge that were consuming lactate by OXPHOS to instead consume glucose by glycolysis, such that MCT1-inhibition has the dual effect of starving the tumour core and increasing the extent of tissue oxygenation (Sonveaux et al. 2008). The latter effect, if confirmed, would make MCT1-inhibition a promising method for radiosensitising symbiotic tumours.

We simulate MCT1-inhibition by CHC—the treatment strategy employed by both Sonveaux et al. (2008) and Busk et al. (2011) *in vivo*—by numerically solving Equations (9)–(12) with the same parameter values from our multi-dimensional parameter space as before, but now with lactate uptake blocked by setting *k_L_* = 0. Contrary to the prediction of Sonveaux et al. (2008), blocking MCT1 has no effect on necrosis in our model system—except when, in cases of high hypoxia, it *decreases* necrosis (Figure 9a–c). These latter cases are mostly, but not entirely, limited to tumours which were non-symbiotic in wildtype, and thus the effect of CHC may be less straightforward than assumed.

**Figure 9:**
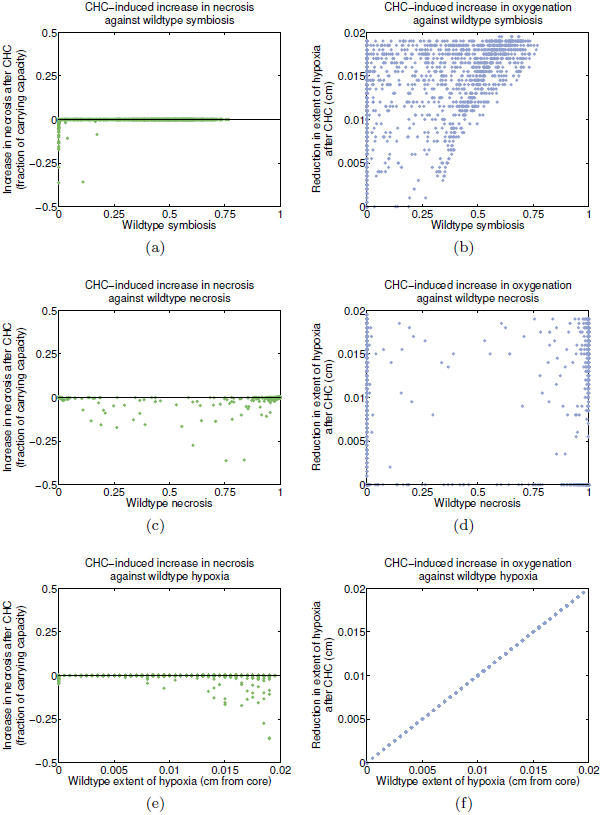
**Effect of MCT1 inhibition on necrosis and hypoxia**. Increase in necrosis (left column) and extent of tissue oxygenation (right column) relative to wildtype, obtained by numerically solving Equations (9)–(12) with 10^4^ parameter points sampled uniformly from the ranges in the bottom portion of Table 1, and comparing these to solutions for the same parameter points but with *k_L_* = 0 to simulate MCT1 inhibition by CHC. Shown are the increase in necrosis and increase in oxygenation as functions of the amount of symbiosis (top row), necrosis (middle row), and hypoxia (bottom row) that were already occurring in the wildtype systems. (c)

CHC treatment does have the expected effect on hypoxia in our model system, in that the extent of oxygenation is increased when lactate uptake is inhibited (Figure 9b–f). However, this effect correlates only loosely with the amount of symbiosis exhibited by the wildtype system, with substantial reductions in the extent of hypoxia occurring even in the absence of wildtype symbiosis. Instead, the effect correlates exactly with the extent of hypoxia that was already present in the wildtype system, with greater reductions occurring for tumours with more extensive wildtype hypoxia.

### 4.4. Experiments suggest weak symbiosis in U87 tumours

One of the model criteria for strongly symbiotic behaviour, as discussed in Sections 4.1 and 4.2, is that the tumour exhibits a strong characteristic preference for lactate over glucose as a metabolic substrate (i.e. that *a* is large). From our *in vitro* experiments on the metabolic characteristics of U87 tumours (outlined in Section 3.3), it is evident that increasing the amount of available lactate leads to a reduction in the extracellular acidification rate (Figure 10a) and a corresponding increase in the oxygen consumption rate (Figure 10b), which together signal consumption of lactate by oxidative phosphorylation. However, there is only a slight decrease in glucose uptake with increasing lactate concentration (Figure 10c), indicating that the cells do not exhibit a strong characteristic preference for lactate but instead consume glucose at a similar rate whether or not lactate is available.

A caveat here is that glutamine and pyruvate were present in the system due to incubation of the cells in standard medium, and, given the interconnectedness of the glucose/lactate/glutamine/pyruvate metabolic pathways, we cannot be sure of the impact of these metabolites. However, we expect our salient result— a qualitative indication of whether or not U87 cells exhibit a marked substrate preference—to hold so long as glutamine and pyruvate do not together suppress a preference that would have been strong in their absence.

Provided this assumption is valid, these experiments suggest that the parameter representing preference for lactate (*a*) is likely to be small, which places U87 glioblastoma tumours approximately into the regime in which only weakly symbiotic behaviour develops—but U87 tumours are highly malignant. This raises the question of how symbiosis might correlate with tumour aggressiveness. It is perhaps possible that symbiotic tumours are stabilised by their metabolic cooperation and insulated from oxygen-dependent treatments, but at the cost of being less malignant than non-symbiotic tumours. This connection, or lack thereof, remains to be explored.

## 5. Discussion

In summary, our aim has been to examine the symbiosis hypothesis put forward by Sonveaux et al. (2008) and address the disagreement in *in vivo* experimental support between their results and those of Busk et al. (2011). We have pursued this by developing a minimal spatial model of dual glucose-lactate consumption, which is an extension, with simplifications, of a non-spatial model presented by Mendoza-Juez et al. (2012). Our main findings from this model are that, while symbiosis arises over a substantial portion of the parameter space, it requires cells to be extremely metabolically active, and whether such a parameter regime is clinically realistic remains to be established. For the most part, the occurrence of symbiosis is mutually exclusive with the occurrence of necrosis in the tumour core, and, as predicted by Sonveaux et al. (2008), the latter is caused by nutrient deprivation (rather than by acidosis or severe anoxia). However, symbiosis and necrosis do not exhibit the same dependence on the model parameters, indicating that symbiosis may not directly ‘rescue’ tumours from nutrient starvation.

We also have found that simulation of MCT1 inhibition does not cause the increase in necrosis in the tumour core that was expected to result from in-creased glucose consumption by cells at the formerly lactate-consuming tumour edge. This places our results in line with those of Busk et al. (2011), who saw no increase in necrosis upon CHC administration *in vivo*. MCT1 inhibition does cause an increase in the extent of tissue oxygenation in our model system, as predicted by Sonveaux et al. (2008), but this correlates with the extent of hypoxia that was present in the wildtype system rather than with its strength of symbiosis. This result gives the promising indication that CHC may be a viable radiosensitisation strategy wherever there is oxidative lactate consumption, independently of whether the tumour exhibits a spatially well-defined symbiosis.

Furthermore, we have demonstrated experimentally that U87 glioma tumours consume lactate but do not exhibit a discernible characteristic preference for lactate over glucose as a metabolic substrate (Figure 10), one of the conditions required for strongly symbiotic behaviour under our modelling framework. U87 tumours are known to be highly malignant, and thus our experimental results compel us to take a step beyond the original hypothesis and raise the new question of whether symbiosis, if indeed it occurs *in vivo*, may be advantageous or disadvantageous (or neither) for clinical malignancy. Under some conditions necrosis appears to be associated with tumour-promoting inflammation and accelerated tumour growth (Degenhardt et al. 2006), such that the relationship between necrosis-reducing symbiotic behaviour and malignancy may not be a simple one.

**Figure 10:**
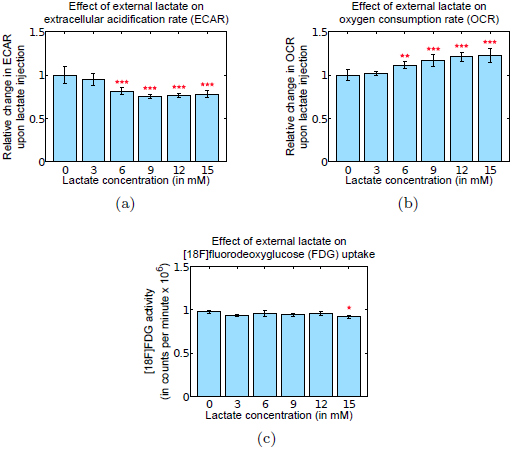
Experimental metabolic dynamics of U87 glioma cells *in vitro*. Metabolic rate dynamics in U87 glioma cells cultured *in vitro* over varying concentrations of lactate in the medium, as outlined in Section 3.3. (a) Extracellular acidification rate (ECAR), a marker of the net effect of glycolysis and lactate consumption; (b) oxygen consumption rate (OCR), a marker of oxidative phosphorylation; and (c) activity of fluorodeoxyglucose ([18F]FDG), a marker of glycolysis. Multiple comparison of a balanced one-way analysis of variance (ANOVA) was performed on the data to test whether the mean measurement for each non-zero lactate concentration was significantly different from the zero lactate concentration. One star indicates a p-value less than 0.05 (but greater than 0.01), two stars indicate a p-value less than 0.01 (but greater than 0.001), and three stars indicate a p-value less than 10*^−^*^4^. Increasing the concentration of available lactate leads to a statistically significant decrease in ECAR and a significant increase in OCR, but a much less significant change in [18F] FDG, suggesting consumption of lactate by oxidative phosphorylation without a strong characteristic preference for lactate over glucose as a metabolic substrate.

We caution that our model is an abstract, minimal representation of tumour glucose-lactate metabolism, and real tumour metabolic systems are a great deal more complicated than what we have considered here. As such, our conclusions should be taken more as theoretical, in-principle statements than as prescriptive of what occurs in real tumours *in vivo*. The framework developed here can be extended in a variety of directions. For example, we do not accommodate vasculature dynamics here, but instead assume the somewhat abstract feature of a fixed capillary shell, and hence the model cannot be used to explore the vasoactive role of lactate (Ido et al. 2003, Hein et al. 2006) or the effect that MCT1 inhibition may have on angiogenesis (Sonveaux et al. 2012). It would also be useful to incorporate intracellular lactate (Parks et al. 2013) into the modelling framework for comparison with experiments by Colen et al. (2011) which induced tumour necrosis by blocking lactate efflux, or to extend the framework further via consideration of glutamate and glutamatergic signalling.

It is worth noting that our tumour radius has been fixed at 0.02 cm throughout, to capture the lengthscale of a typical avascular tumour, and for simplicity we have not considered varying this here. However, we expect that the consequences would be straightforward—a smaller tumour would exhibit fewer spatial features due to less-pronounced gradients, and a larger tumour would develop a necrotic core with a surviving region, of a length similar to that considered here, in which the metabolic dynamics predicted by our model would occur. A systematic examination of different lengthscales, however, could prove useful for determining the earliest point in tumour growth at which spatial features may develop. Similarly, we have asserted that our model system can be considered to describe localised pockets within a large, heterogeneous tumour; but it would be beneficial to explicitly model such a tumour as a whole, as this would introduce spatial complexities and facilitate more direct comparisons with medical neuro-imaging *in vivo*.

Our *in vitro* experiments were preliminary, and served simply to facilitate some speculation as to whether tumours might exhibit some characteristic preferences regarding metabolic substrates, and where such preferences might place the tumours in our model parameter space. These experiments should in future be extended to a series of microfluidic (Whitesides 2006, Huang et al. 2011, Zhang & Nagrath 2013) or *in vivo* measurements of metabolic consumption for a variety of tumour types—similarly to Sonveaux et al. (2008) and Busk et al. (2011) but with local glucose and lactate consumption rates measured quantitatively—as these would help to place tumours in our model parameter space far more accurately than we have attempted here. Such experiments could thereby more conclusively establish whether symbiosis is likely to be a significant feature of tumours *in vivo*.

Nevertheless, the minimal spatial model presented herein has enabled some predictions which can be tested experimentally—in particular, the relationship (or lack thereof) between the occurrence of symbiosis and degree of clinical malignancy can be investigated—and which support the *in vivo* observations of Busk et al. (2011) with regard to CHC-induced necrosis over those of Sonveaux et al. (2008). Hence, we hope that this work is illustrative of the gains that can be made by integrating mathematical modelling with experiments for clearer and more rigorous testing of hypotheses in oncology.

## Acknowledgements

JBM and PKM acknowledge the generous hospitality of the Instituto de Matemática Aplicada a la Ciencia y la Ingeniería, Universidad de Castilla-La Mancha. JBM would like to thank Dr. Gregory A. Ross for helpful feedback on the manuscript.

